# The Role of Migration in the Evolution of Phenotypic Switching

**DOI:** 10.1101/003442

**Authors:** Oana Carja, Robert E. Furrow, Marcus W. Feldman

## Abstract

Stochastic switching is an example of phenotypic bet-hedging, where an individual can switch between different phenotypic states in a fluctuating environment. Although the evolution of stochastic switching has been studied when the environment varies temporally, there has been little theoretical work on the evolution of phenotypic switching under *both* spatially and temporally fluctuating selection pressures. Here we use a population genetic model to explore the interaction of temporal and spatial variation in the evolutionary dynamics of phenotypic switching. We find that spatial variation in selection is important; when selection pressures are similar across space, migration can decrease the rate of switching, but when selection pressures differ spatially, increasing migration between demes can facilitate the evolution of higher rates of switching. These results may help explain the diverse array of non-genetic contributions to phenotypic variability and phenotypic inheritance observed in both wild and experimental populations.

## Introduction

In a static environment, phenotypes might be expected to evolve towards some optimum, with selection ultimately producing an adapted population with minimal phenotypic variation. However, when selection varies through time and space, mechanisms that maintain diversity of phenotypes may evolve. Although individuals experience no immediate fitness benefit, these mechanisms could act as a form of bet-hedging, increasing the long-term survival and growth of a lineage [1, 2]. One such mechanism involves mutation that produces variation among two or more phenotypes. This is known as phenotypic or stochastic switching, and has been observed in a diversity of organisms such as viruses [3], yeast [4–6], and bacteria [7–9].

Stochastic switching can describe multiple stable expression states for a gene or genetic pathway. These multiple states may correspond to differences in epigenetic marks (mammalian examples reviewed by Daxinger and Whitelaw [10], plant examples reviewed by Henderson and Jacobsen [11]), or result from positive-feedback transcriptional loops such as the galactose-signalling network in the yeast *Saccharomyces cerevisiae* [4] or DNA uptake pathways in the soil bacterium *Bacillus subtilis* [8]. Genetically determined variation can allow offspring to be phenotypically similar to parents, while phenotypic plasticity may cause phenotypes to differ greatly within a genetic lineage. Stochastic switching can produce phenotypic variability with familial correlations intermediate between these two extremes (as is seen in the contributions of DNA methylation variation to heritability of phenotypes in the plant *Arabidopsis thaliana* [12]).

Theoretical studies have found that these intermediate phenotypic correlations should evolve in tune with the correlation between environments of parent and offspring. Early studies [13, 14] found that when the environment fluctuates periodically between two states with different optimal phenotypes, the switching rate between phenotypic states should evolve to approximately 1*/n*, where *n* is the number of generations between temporal environmental changes. In this way, the switching rate matches the parent-offspring phenotypic correlation to the correlation in environment between generations. However, Salathé *et al.* [15] showed that evolutionarily stable switching rates became close to zero as the variability in temporal fluctuations increased, or if fitness costs were asymmetric between the two environments. Including the potential for non-random phenotypic switching, Kussell and Leibler [16] found that costly sensing was favored in rapidly changing environments, while pure stochastic switching was favored in slowly changing environments.

Although natural populations might be expected to experience variability through both space and time, there are few studies of stochastic switching in which the environment varies both temporally and spatially. Arnoldini *et al.* [17] studied the evolution of sensed switching in a population with multiple spatial patches and found that the relative balance of sensed and stochastic switching depended on the accuracy of the environmental stress signal. Unlike the model we present here, that study did not explore different migration rates, nor did it allow for temporal environmental correlation. From her analysis of non-inherited phenotypic variability in a spatially and temporally varying environment, Moran [18] argued that the optimal level of variability is zero. However, in the case of stochastic switching, parents and offspring will generally have non-zero correlation in phenotype: a scenario that is not possible in Moran’s model.

We expect the evolution of stochastic switching to be strongly influenced by spatial heterogeneity; in its absence a temporal change in the environment is experienced by the entire population. On the other hand, in a spatially heterogeneous environment, migrants may experience a new environment in which they compete with residents that have a high frequency of the phenotype that is optimal for that deme. If switching interferes with this local adaptation, it may not evolve even when there are high rates of migration between demes. But if higher rates of switching are extremely beneficial to recent migrants, a greater rate of dispersal may select for more switching. A recent theoretical analysis, focusing on the mathematically tractable case with strict symmetry of selection and constant waiting times before environmental change, demonstrated that migration can supplant the need for switching [19]. However, such stringent symmetry conditions may characterize only a small subset of the ecological scenarios in which switching can be adaptive.

In this paper, we study the evolution of stochastic switching in a population that is spatially subdivided into demes, with a range of selection regimes, and where each deme also experiences stochastic temporal variability in selection. By exploring the invasion of different switching rates across a range of migration rates and temporal and spatial selection regimes, we analyze the relative roles of spatial and temporal fitness variability in determining the evolutionarily stable rates of stochastic switching. We find that the dynamics of this system are determined by a complex interaction between migration and stochastic temporal fluctuations. When different temporal states produce similar strengths of selection, increased migration selects for lower rates of switching or has a minimal effect, depending on the fitness regime. Unlike previous results, we find that when some temporal states exert stronger selection than others, increased migration can select for higher rates of stochastic switching. This surprising finding highlights the interaction between spatial and temporal environmental variability in determining the evolution of phenotypic switching.

## Model

We use an explicit population genetic model, tracking the allele frequencies at a modifier gene that determines the rate of stochastic switching in a spatially heterogeneous metapopulation. Our goals are to explore the conditions under which this switching can evolve when fitness varies across both space and time, and to understand how the evolutionary dynamics in this model differ from those in models that allow only temporal variation in fitness.

The population is spatially divided into two demes, *E_x_* and *E_y_*, each of which is effectively infinite in size. Each individual in the population is haploid and defined by two biallelic loci: a major locus *A/a* which controls the phenotype and thus the fitness of the individual, and a modifier locus *M/m* which controls the phenotypic switching rate between phenotypes *A* and *a*. Switching occurs only at the phenotypic locus, at a rate that is assumed to be the same in both directions. A possible mechanism for switching is epigenetic control of gene expression through variation in levels of methylation or chromatin loop formation. Therefore, the *M/m* modifier locus can be interpreted as a genetic locus that influences the transition between two different stable expression levels of an allele. Examples of such loci are the *DNMT* genes, which have been shown to have a role in the establishment and regulation of cytosine methylation [20]. Because this locus may be genetic or epigenetic, we explore a broad range of switching rates.

Within each deme the environment varies temporally between two states, *T*_1_ and *T*_2_. To incorporate random temporal variation, the waiting times between environmental changes are drawn from a gamma distribution. As a measure of environmental variability we use the parameter *ψ*, defined as the variance of the gamma distribution divided by the square of its mean. This allows us to test a range of distributions between pure periodicity (*ψ* = 0) and an exponential waiting time (*ψ* = 1) by fixing the mean of the distribution while varying the variance.

Selection acts only on the phenotypes *A* and *a*, and the fitness of these two phenotypes is determined by the spatial and temporal states which they inhabit. The modifier locus *M/m* is assumed to be selectively neutral. At each time-step, individuals first experience selection, followed by switching. Blanquart and Gandon [21] have demonstrated that recombination rates may play an important evolutionary role in models with migration; we assume recombination occurs between the modifier locus and the phenotype locus at rate *r*. After recombination, the offspring can migrate at equal rates between the two demes. The recursions representing the change in two-locus genotype frequencies at every generation are presented in the **Supplementary Material**. Because selection is local, with individuals only competing within their deme, the environmental state cannot be interpreted as another genetic locus or phenotypic state.

For simplicity, we assume that, within each deme, phenotype *A* is favored in one temporal state and phenotype *a* is favored in the other. Temporal states are denoted by *T*_1_ and *T*_2_. The environment in which allele *A* is optimal is *T*_1_ in deme *E_x_* and *T*_2_ in deme *E_y_* . Under this assumption, the fitness regime can be represented by the four different selection coefficients in **Table 1**. After selection, alleles *A* and *a* switch to the opposite state at rate *µ_M_* or *µ_m_*, if they are on *M* or *m* genetic backgrounds, respectively. Note that because *ψ* is generally non-zero, the two demes will experience independent changes in temporal state. In these cases, the condition that selective forces are opposite in the two demes is not as restrictive as it seems; there are simply times when selection within a deme favors *A*, while at other times it favors *a*.

**Table 1.** Fitnesses for each of the combinations of phenotype (*A/a*), deme (*E_x_/E_y)_*, and environmental state (*T*_1_*/T*_2_).

| Deme | Temporal State | Phenotype | Fitness |
| --- | --- | --- | --- |
| $E_x$ | $T_1$ | $A$ | 1 |
| | | $a$ | $1 - s_1$ |
| | $T_2$ | $A$ | $1 - s_2$ |
| | | $a$ | 1 |
| $E_y$ | $T_1$ | $A$ | $1 - s_3$ |
| | | $a$ | 1 |
| | $T_2$ | $A$ | 1 |
| | | $a$ | $1 - s_4$ |

### Description of the simulation

The population is initiated with only the *M* allele at the modifier locus. After allowing the population to evolve for 1000 generations, we introduce a small fraction (10^*−*4^) of allele *m*, with a new switching rate, into the population. Evolution then proceeds for 5000 generations after which we evaluate whether or not the new modifier invaded; invasion is declared if, at the end of this invasion trial, allele *m* has a frequency strictly larger than its initial 10^*−*4^. In this case, we expect *m* to have had a selective advantage over *M*, and that it would reach fixation if we allowed the population to continue evolving (see also **Supplementary Figure S1**).

To find the evolutionarily stable switching rate, we repeat this invasion trial 500 times, or until no new modifier is able to invade for 20 consecutive iterations. We start a simulation run with a randomly chosen value of the (wild-type) switching rate for the *M* allele. The switching rate corresponding to *m* is chosen as the product of this wild-type rate and a random number generated from an exponential distribution with mean 1. After each iteration, if *m* invades, it becomes the new resident allele in the next invasion trial. The final switching rate after these 500 trials is considered to be the evolutionary stable switching rate.

Because of the stochasticity introduced by the *ψ* parameter, the final stable switching rate is computed as the average of the stable switching rates obtained in 10 different runs of the simulation presented above.

### Migration can decrease the stable switching rate

We first explore the dynamics of the system in a symmetric fitness regime, assuming both spatial and temporal symmetry in the selection pressures (*s*_1_ = *s*_2_ = *s*_3_ = *s*_4_). **Figure 1** shows the results when the expected time before an environmental change is 10 generations in both demes and this temporal change is sampled from a gamma distribution with variability parameter *ψ* represented by line color.

**Figure 1.**
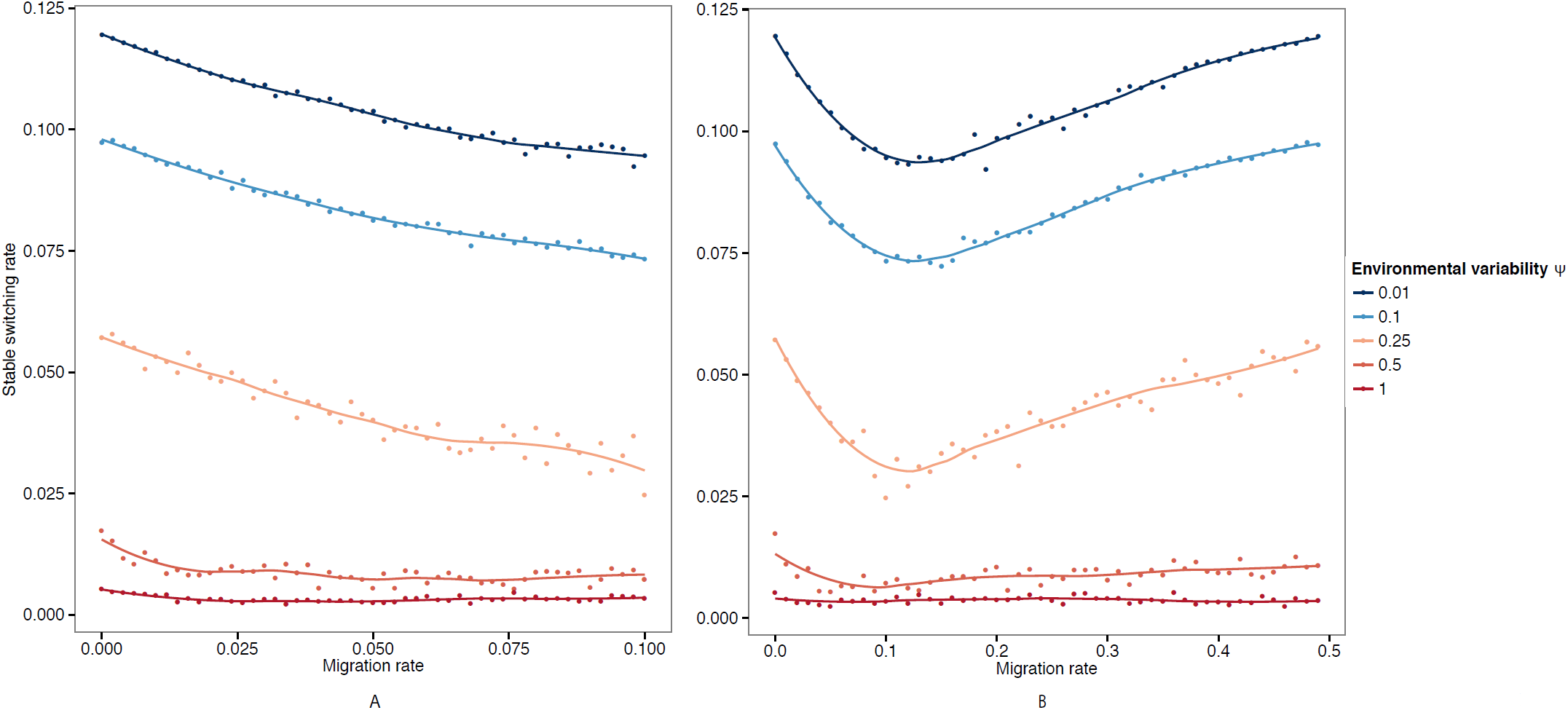
The stable switching rate when selection is symmetric in space and time. All selection coefficients are 0.1. Recombination rate is 0.1. The expected time before an environmental change is 10 generations in both demes, and the colors correspond to different levels of environmental variability *ψ*. Environmental variability *ψ* = 0 corresponds to the exactly periodic case of an environmental switch every 10 generations. Environmental variability *ψ* = 1 represents the case where both environments change with an exponential waiting time of 10 generations. **Panel A**: small migration rates; **Panel B**: larger range of migration. The plotted curves represent a fit to the data using a generalized additive model with penalized cubic regression splines.

**Figure 1, Panel A** presents the evolutionarily stable switching rate as a function of small increases in the migration rate between demes. With no migration, we recapture the results from models on the evolution of phenotypic switching when there is only temporal heterogeneity in the system. In this case the evolutionarily stable switching rate is on the order of 1*/n*, where *n* is the mean waiting time before an environmental change [13–15, 22]. Increasing the variability parameter *ψ* (by increasing the variation in waiting time) decreases the evolutionarily stable switching rate, which can then be orders of magnitude less than 1*/n*, consistent with previous results in the absence of spatial heterogeneity [15]. As the migration rate increases, the evolutionarily stable switching rate decreases. This result may stem from the increased heterogeneity in selection that any particular lineage may experience, as occasional migration simulates variability in waiting time before an environmental change.

**Figure 1, Panel B** shows the stable switching rate for a larger range of migration rates, between 0 and 0.5. The observed initial decrease in the stable switching rate is reversed for increasing migration rates. This result is intuitive; when the population is well mixed (migration rates equal to one half), the stable switching rate should be the similar to the case of no spatial heterogeneity (migration rates equal to zero).

The results shown in **Figure 1** are invariant to the strength of symmetric selection pressure used (**Supplementary Figure S2**), and the qualitative pattern is robust to different environmental mean waiting times, both when the times are the same in the two demes (**Supplementary Figure S3, Panel A**), and when they differ between demes (**Supplementary Figure S3, Panel B**).

Moreover, this dip in switching rates as migration increases is robust to asymmetry in the overall strength of selection between the two demes. **Supplementary Figure S4** shows results where the fitness reduction of the maladapted phenotype is larger in one deme than the other, *s*_1_ = *s*_2_ *> s*_3_ = *s*_4_. We expect the evolutionary dynamics within the former deme to dominate the system and, as observed in the symmetric cases, environmental variability reduces the stable switching rate (**Supplementary Figure S4**).

### Migration can increase the stable switching rate

When selection favors one phenotype more strongly in the first deme, and the opposite phenotype in the second deme, higher rates of migration can lead to the evolution of higher switching rates. In this case, there is asymmetry in selection both within and between demes (*s*_1_ *> s*_2_, *s*_3_ *> s*_4_). **Figure 2, Panel A** illustrates the difference between the symmetric regime presented above (all selection coefficients equal) and this regime in which higher migration rates select for higher switching rates. As the level of asymmetry in selection within demes increases, the curves change from dipping to monotonically increasing with migration. We expect that selection may often be asymmetric in strength within a deme, so this finding greatly expands the range of selective regimes that might allow the evolution of switching. The source of this effect may be the distribution of phenotypes within a deme; when seletion is asymmetric within a deme, certain generations will have one phenotype dominating that subpopulation. If there is high migration into a deme with opposing selective forces, most migrants will carry the same phenotype, and switching might be very beneficial as a means to compete within a population that is already much better adapted.

**Figure 2.**
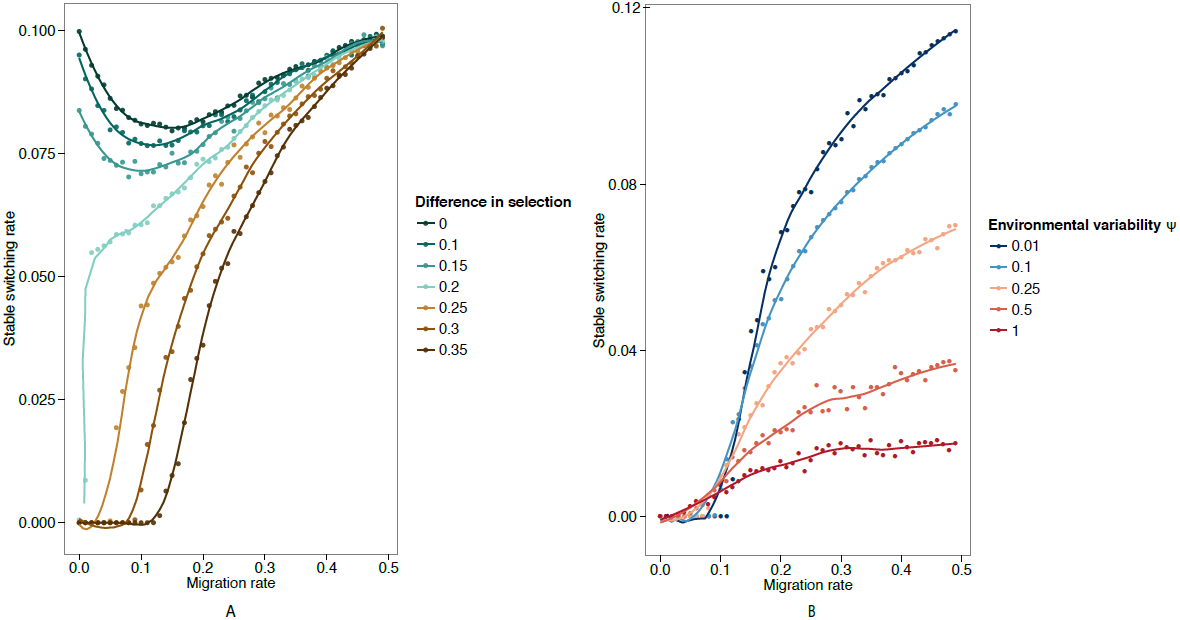
The stable switching rate when selection strengths vary temporally. **Panel A**: The selection coefficients are *s*_1_ = *s*_3_ = 0.4, and *s*_2_ = *s*_4_ with the difference in selection pressure as presented in the legend. Recombination rate is 0.1. The expected environmental period is 10 generations in the both demes. The environmental variability parameter *ψ* is equal to 0.1. **Panel B**: The selection coefficients are *s*_1_ = *s*_3_ = 0.4, and *s*_2_ = *s*_4_ = 0.1. Recombination rate is 0.1. The expected environmental period is 10 generations in the both demes, and the colors correspond to different levels of environmental variability *ψ*. Environmental variability *ψ* = 0 corresponds to the deterministic case of an environmental switch every 10 generations. Environmental variability *ψ* = 1 represents the case where both environments change with an exponential waiting time of 10 generations. The plotted curves represent a fit to the data using a generalized additive model with penalized cubic regression splines.

**Figure 2, Panel B** focuses on the case where this difference in selection is equal to (*s*_1_ = *s*_3_ = 0.4, *s*_2_ = *s*_4_ = 0.1). Similar to the results observed in **Figure 1**, higher environmental variability *ψ* generally selects for lower switching rates.

These qualitative results are robust to different expected waiting times before temporal environmental changes (**Supplementary Figure S5, Panel A**), as well as to differences in expected waiting time between the two demes (**Supplementary Figure S5, Panel B**). As migration rates increase, the evolutionarily stable switching rate approaches the rate for a heterogeneous population: approximately the inverse of the expected waiting time (**Supplementary Figure S5, Panel A**).

### Recombination decreases the stable switching rate

The evolutionarily stable switching rate declines linearly with increasing recombination rate *r* for all of the different selection regimes presented above. **Supplementary Figure S6** presents a case in which there is symmetric selection (selection coefficients are all 0.1), the environments switch with an expected period of 5 generations in both demes and the migration rate *ν* is 0.05. The stable switching rate decreases linearly with increasing recombination rate. This simple result suggests recombination may partially supplant the need for switching. From the perspective of a modifier allele, a recombination event can simulate the act of switching, as that allele may end up on a background with a different phenotype.

## Discussion

Organisms experience environmental heterogeneity through both space and time, and their descendants may experience different environments due to migration and temporal environmental changes. Here we explore the evolution of stochastic switching between different phenotypic states in such a heterogenous environment. A focal parameter of our analysis is the rate of migration between demes, which interacts with spatial and temporal environmental heterogeneity in selection to affect the long-term growth rate of a lineage. With all else held equal, higher migration rates correspond to lower correlations between the demes of parents and their offspring, and therefore a greater spatial contribution to variation in fitness.

When migration between demes is relatively infrequent, our model reiterates the message of previous theoretical models; stochastic switching can evolve, and the stable rates of switching should approximate the inverse of the expected number of generations before an environmental change. However, when migration rates are larger, lineages experience additional spatial variation in selection. This results in two qualitatively different possibilities: spatial variation may reduce switching for small increases in migration rates but have minimal effect as migration becomes very frequent, or greater spatial variation in fitness may induce selection for higher rates of switching. To understand the ecological implications of this, we consider the conditions that separate these qualitative regimes.

Without temporal asymmetry in selection, the qualitative role of spatial heterogeneity depends on the degree of migration in the metapopulation. For small amounts of migration, switching is reduced, because migration produces heterogeneity in the selection experienced by a lineage, and may partially supplant switching as a way to match phenotype to environment. For higher migration, the metapopulation is essentially a single population, and the results mirror those found for single population models. The relevant scale of migration will depend on the spatial scale at which selection varies in the environment which they inhabit.

When the strength of selection in the two demes is asymmetric, migration can select for higher switching rates at evolutionary equilibrium. One possible explanation for this is that a migrant may move to a highly disadvantageous novel environment, where the benefit of switching to the new optimal phenotype outweighs the risk of switching at the wrong time. This is consistent with the idea of bet-hedging as a protection against occasional highly stressful events [23, 24]. One example could be seed dormancy as a bet-hedging mechanism in annual plants [25]. We did not study the evolution of migration rates in our model, but it could be interesting to determine conditions under which migration and switching evolve in concert or in opposition, since they both influence the variation in fitness that a lineage will experience.

Although previous theoretical work on switching has been framed as relevant to bacteria, viruses, and yeast, epigenetic mechanisms in plants may be modeled in a similar manner. For example, in a clonal line of dandelions Verhoeven *et al.* [26] found that a variety of DNA methylation changes could be induced by simulating a range of environmental stresses such as herbivory or high environmental salt concentrations – such stresses could vary through both space and time in a natural population. Many of these epigenetic changes were transmitted faithfully for several generations. For a population in which all individuals have the same probability of experiencing such a stress, then a model of stochastic switching could represent the dynamics of DNA methylation across generations. Despite the fact that these epigenetic changes stem from an environmental cue, randomly occurring cues or variable responses to a cue may effectively produce stochastic switching between phenotypic states. More work is needed to illuminate the effects of these epigenetic states on fitness and thus on the evolution of stochastic switching.

Here we show that migration does not have the same effect in all ecological scenarios. In some cases it can supersede stochastic switching by allowing migrants to avoid temporal environmental changes. In other cases migrants may be exposed to stressful environments, producing selection for high rates of switching. Care must be taken when considering the adaptive role of stochastic switching in a natural population. Does the population experience occasional strong selection, or does the environment cycle through a variety of mild selective events? Is there spatial heterogeneity in fitness, or do temporal environmental changes dominate? For example, with the looming challenge of antibiotic resistance among yeast and bacteria, we might want to consider how drug choice, treatment timing, and potential microbial migration between human hosts can interact to select for higher or lower mutation rates [27]. Under spatial environmental variation, conditions for the evolution of switching mechanisms may not be as restrictive as previously thought [15, 28]. These results offer insight into the occurrence of high levels of phenotypic variability in many populations, and call for research on switching, epigenetic inheritance, and mutation rates that explicitly considers spatial heterogeneity.

## Acknowledgments

We thank members of the Feldman laboratory for helpful discussion.

